# Predication of the Post Mining Land Use Based on Random Forest and DBSCAN

**DOI:** 10.1101/2023.05.30.542990

**Authors:** Qiang Bo, Pinhan Lv, Ziguan Wang, Qian Wang, Zechuan Li

## Abstract

Mine reclamation is one of the most important stages of the mining activities in line with the basic principles of sustainable development. In this study, different post-mining land uses are evaluated in the Hongliulin mining area, which is located in Shen mu country of China. 145 soil samples were collected in the May,2021 by using the soil auger, and the sampling depths were 0-20 cm. The sampling points contains 45 to be reclaimed samples and 100 existing classification land use types. 14 environmental factors including soil organic matter (SOM), total nitrogen (TN) and other soil nutrients and terrain factors were extracted and calculated based on laboratory test and digital elevation map. Meanwhile, random forest classier was utilized to determine the post-mining land use based on GINI index and 14 environmental factors by using 100 existing classification land use types. 82 of 100 samples were utilized to build the model and the other 18 samples were utilized to validate the accuracy of the classification model. Density-Based Spatial Clustering of Applications with Noise (DBSCAN) techniques were utilized to determine the specific land use types in cropland, shrub and grassland. the random forest classier showed a great prediction capability, with only 1 miss-classified sample in the validation data-set, the accuracy of the classification model was 95%. Among the 45 samples to be reclaimed, 15 samples were classified into cropland, 23 samples were classified into shrub and grassland, 5 samples were classified into arbor land, 2 samples were classified into solar station. By using DBSCAN, the 58 samples classified into cropland in terms of the post-mining were separated into cluster 1 and 2, cluster 1 comprises 24 samples while cluster 2 comprises 34 samples. Among the 61 samples classified into shrubland and grassland, Cluster -1 comprises 24 samples and Cluster 0 comprise 22 samples. The results could provide technological support for land reclamation determination. The content of TN of C1 is 5 times more than C2 and 4 times more than C3. Also, the K valve of C1 column is maximum and over 0.4, which means the soil particle is relatively smaller and the soil texture of it is sandy loam. The TWI of C1 is minimum, in consideration of a positive valve of profile curvature and a negative valve of plan curvature. In terms of the 45 to be reclaimed samples, 15 samples were classified into C1, 23 samples were classified into C2, 5 samples were classified into C3, 2 samples were classified into C4. The valve of K and content of soil nutrients of the samples classified to be C1 column(C1-C) is maximum. Simultaneously, the slope steepness(°) is below 5° and is perceived as gentle surface. Cluster 1 comprises 24 samples and the average content of TN, AP and SOM is 0.566g/kg, 11.93mg/kg and 19.975g/kg respectively, while Cluster 2 comprises 34 samples and the average content of TN, AP and SOM is 0.304g/kg, 3.12mg/kg and 8.36g/kg respectively. The result may contribute to the land use planning and idle land utilization strategy.

## 1. Introduction

Mining activities have brought many negative impacts on land, economy. The contradiction between man and land has become increasingly acute. At present, about 30% mining areas in China have entered the middle-aged and old age after a long period of mining. If the land resource cannot be used effectively, it will cause a serious waste of land resources, and reclamation is an effective way to protect land resources [1]. At the same time, it can also improve the land utilization rate in the mining area, which is of great significance for adjusting the economic structure of the mining area and changing the growth mode [2]. As the arable land area has been approaching the “alarming line” in recent years, the land use types in mining areas are still extensive [3].

Land reclmation evaluation required both soil nutriments index and topographic index [4]. In the process of establishing an evaluation model suitable for the mining area, the first thing to consider is the classification of the land use types of the land to be reclaimed. There are four main types of land use in the mining area in China, including cultivated land, shrub land, grassland, arbor land and solar plant [5]. In the case of giving priority to reclamation as arable land, the soil fertility and arability of the area should be evaluated [6]. Specifically, the emergence of a large amount of ground fissures and land deformation caused by mining activities will lead to thinning of the soil layer, destruction of the soil structure, loss of soil fertility and other phenomena that are not conducive to soil development [7]. With the development of ground fissures, it is bound to erode the fertile soil layer. It causes a large amount of soil organic matter and nutrient loss, deterioration of soil physical and chemical properties, soil compaction, soil quality deterioration, soil ventilation and water permeability reduction, and rapid decline in soil fertility and quality [8]. Therefore, the impact of soil nutrients should be fully considered when evaluating the suitability of reclamation. If the land cannot be restored to cultivated land due to suitability constraints, its suitability for reclamation into forest land shall be evaluated according to its site conditions. Finally, the suitability of reclamation as a solar power plant is evaluated based on factors such as terrain undulation and terrain fragmentation. Secondly, after determining the direction of land use, according to the classification of cultivated land, shrubs and grassland, its suitability grade should be divided. Select the corresponding reclamation scheme according to different suitability grades.

Soil organic matter (SOM) and total nitrogen (TN) was perceived as the most essential nutrients in reclamation process. Soil organic matter is the main source of trace elements such as nitrogen, phosphorus, and sulfur required by plants. According to statistics, more than 75% of the nitrogen in the topsoil (0-20cm) of the study area exists in the form of organic state. The mineralization of soil organic matter further promotes the degradation and transformation of microorganisms. During the process of degradation and transformation, microorganisms can adjust the data and release nutrients [9,10]. The structure of soil will be affected by soil organic matter. During the extensive research on sandy soil, it is not difficult to see that organic matter can effectively increase the agglomeration effect of soil particles, making it softer, structured and sticky soil, increasing the tillability of the soil. Nitrogen is an important element of chlorophyll in plants. In addition, nitrogen also plays an important role in the formation of proteins and the pairing of nucleic acids, and can help chlorophyll in the conversion of inorganic matter to organic matter and the conversion of light energy and chemical energy [1]. Conversion, the product formed by chloroplasts in photosynthesis is glucose. Meanwhile, the available nutrients such as available phosphorus was crucial for plants and vegetation growth. Thus, it was necessary to evaluate the SOM, TN and other available nutrients in the reclamation process.

Multi-criteria decision-making (MCDM) has been widely utilized to evaluate the suitably of land reclamation, among these studies, PROMETHEE and TOPSIS was proved to be effective strategy for classification of land use types in terms of reclamation [11]. However, the complexity in the classification of land use type is generally non-linear, thus the MCDM may fail in the accurate classification of land use types as MCDM was only susceptible to linear classification. Machine learning has shown superiority in dealing with the classification of non-linear, thus, it was essential to make post-mining land use classification by using machine learning [1].

Taking the Hongliulin mining area as an example, about 30km^2^ of golfs have appeared, and the land is idle. In addition, more working faces are being mined one after another, and a large number of abandoned land will appear [8]. Several severe devastations to the land resource was brought to the land due to the mining activities. 1). About 1/4 area has been proved to have a ground fracture/collapse pit or other forms of subside, which cause the transportation such as bridge and country road stalled or nearly paralyzed. 2). Plentiful cropland has been occupied to serve the mining. Once mining is complete, the coal gangue and the pipeline as well as other facility was deserted, which caused idle land and abandoned land. 3). As is demonstrated in the previous research, the content of soil nutrient, comprise total nitrogen, available phosphorus, available potassium and soil organic matter decrease with the mining activities, which cause the reduction of output. The major aims of this manuscript was to: 1). extract environmental factors affecting post-mining land use classification. 2). build the post-mining land use prediction model based on random forest. 3). decide the specific land use types in cropland and shrubland & grassland by using DBSCAN clustering method.

## 2. Study Area

The area is located in the northern part of the Loess Plateau in northern Shaanxi, on the southern edge of the Mu Us Desert (Figure 1). The western part is a wavy dune land, and the eastern part is a loess geological valley [1]. The study area belongs to the first panel of the Red willow mining area. The terrain is open and the loess gully develops. The terrain is generally high in the northwest, low in the southeast, high in the middle, and low in the north and south. The study area belongs to the mid-temperate semi-arid continental climate, with cold winters and hot summers. The temperature difference between day and night is with huge disparity, about 20°C maximumly. From November to March of the following year, the maximum depth of frozen soil is 146cm; the maximum snow thickness is 12cm; the monsoon period from the beginning of January to the beginning of May, mostly northwest wind, the average annual wind speed is 2.5m/s, and the maximum wind speed is 25m/ s, the annual average temperature is 8.5 °C, the extreme maximum temperature is 38.9 °C, the extreme minimum temperature is −28.5 °C, the annual average precipitation is 434.10 mm, and the concentration is mostly concentrated in 7, 8 and 9 months; the annual average evaporation is 1907.2 to 2122.7 mm, about 4 to 5 times the amount of precipitation [8].

**Figure 1.**
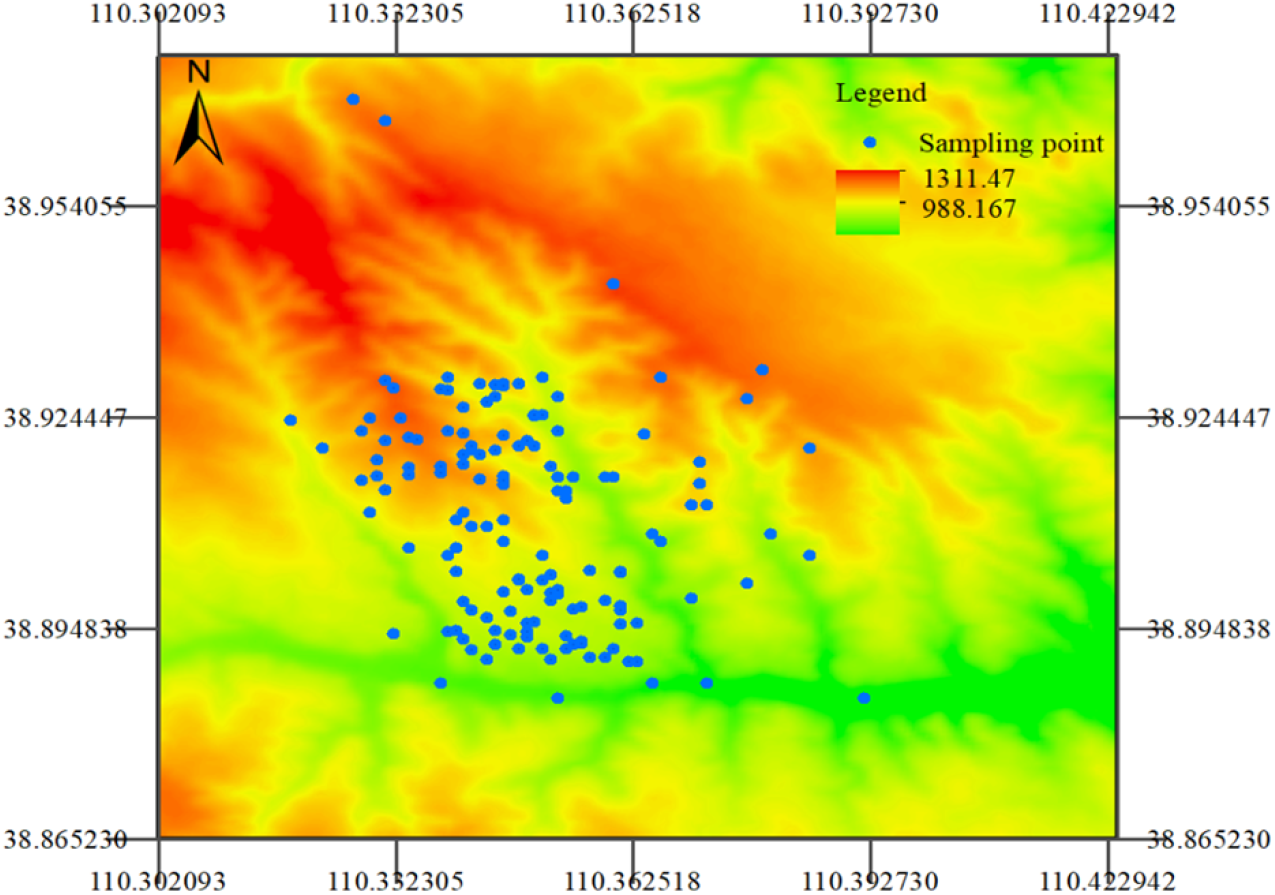
Study area and sampling points.

## 3. Material and Method

145 soil samples were collected in the May 2021 by using the soil auger, and the sampling depths were 0-20 cm. For each sample, soil from 5 points within a 50 m radius area was taken and put together to make a mixed sample for soil chemical and physical analysis. The sampling points contains 45 to be reclaimed and 100 existing classification land use types. The sampling soil type is aeolian sandy soil.

The samples comprise 4 types of previous land use and now are all idle and destructive due to the mining activity (Table 1). Cropland and shrub land occupy 81% of the total land use types. Among the cropland, 85% of which is cornfield, the other is murphy. The shrub and grass land comprise salix mongolica, Artemisia desertorum, caragana microphylla and Astraglus adsurgens. The features which were utilized to establish the classifier comprise terrain factors and soil nutrients.

**Table 1.**
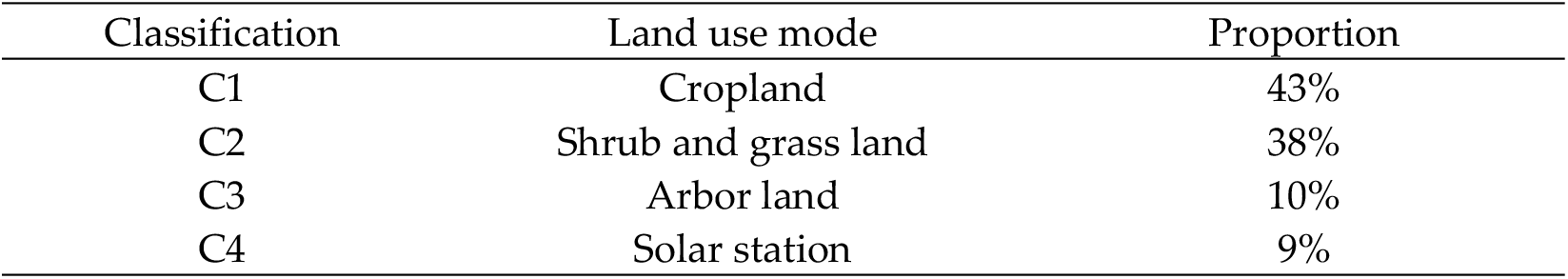
Current land use type of 100 samples.

| Classification | Land use mode | Proportion |
| --- | --- | --- |
| C1 | Cropland | 43% |
| C2 | Shrub and grass land | 38% |
| C3 | Arbor land | 10% |
| C4 | Solar station | 9% |

### Terrain factors

1) K factor. K factor is one factor embodied in the USLE model and is utilized to evaluate the erodibility of the soil, the formula of the K factor is as below [2]:

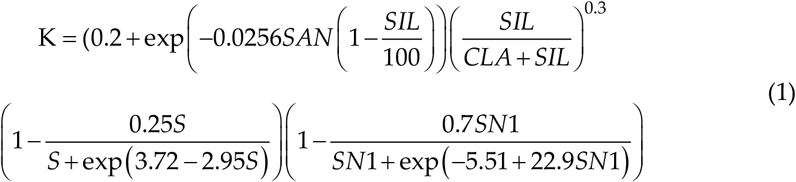

Where,

SAN, SIL, CLA, S refer to the percentages of sand, silt, clay and SOM in the soil (%), respectively. SN1=1-SAN/100.

Generally, the K valve of sandy loam is higher than the loamy sand due to the particle of it is smaller. The K factor limited an area to be reclaimed for cropland as the large particle will result in the nutrients loss.

2) Topographic Wetness Index. TW index is considered as one quantitative and accurate description of soil moisture content in a small watershed, which proved to be one major limiting factor for agriculture and reclamation in the arid and semiarid area. TWI is calculated as below [1]:

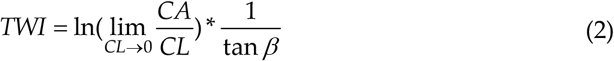

Where,

CA refers to the flow accumulation and CL refers to the flow direction. β refers to the slope steepness(°). A high valve of TWI usually signifies a larger flow confluence area, which lead to a higher soil moisture content and ability to generate runoff on account of more susceptible to achieve saturation.

1. Curvature and NDVI. Generally, profile curvature is parallel to the slope and indicate the direction of the maximum slope. It affects the acceleration and deceleration of the flow through a surface. Plane curvature is a direction perpendicular to the maximum slope. It affects the convergence and dispersion of flows through a surface. A negative value of profile curvature indicates that the surface of the cell is convex upward and the flow rate will decrease. A positive valve indicates that the surface opening is concave upwards and the flow rate will increase. A value of zero indicates that the surface is linear. A positive valve of plan curvature will separate the runoff will a negative one will accumulate the runoff. the Normalized Difference Vegetation Index (NDVI) can reflect vegetation coverage, which is generally high in the arbor land and cropland in summer [8].

Soil nutrients are taken to evaluate the fundamental property of the post mining land. The analyses items included soil organic matter (SOM), total Nitrogen (TN), available Phosphene (AP) and available Kalium (AK). SOM was measured with the K2Cr2O7 heating method [1]; TN, with Semi-micro Kjeldahl method [2]; AP, with the alkaline hydrolysis NaHCO3-extraction-Mo-Sb-Vc-colormetry [8]; and AK, with ammonium acetate extracts Flame photometer [1]. The soil texture was measured with the soil sieve. The relative deviation is less than 5%. All of the above components were analyzed at the laboratory of the Chinese Academy of Agricultural Sciences. All of the terrain factors are finished in the Arcgis 10.1 by using a 5-meters digital elevation map.

### 3.1 Mathematic Method

#### 3.1.1. Random Forest Classification

Random forest classification is an ensemble learning firstly proposed by Breiman [12] and Adele Cutler which processes a classifier that utilizes multiple trees to train and predict samples[12]. Simply put, a random forest is made up of multiple CART (Classification and Regression Tree). For each tree, the training set they use is back-sampled from the total training set, which means that some samples in the total training set may appear in the training set of a tree multiple times, or may never appeared in the training set of a tree. When training the nodes of each tree, the features used are randomly extracted from all the features according to a certain proportion. According to Leo Breiman’s suggestion, the total number of features is assumed to be M. This ratio can be sqrt (M), 1/2sqrt(M), 2sqrt(M).

A tree is made up of all of the features (soil nutrients and terrain factors) based on the purity of mathematics, which means target variables must be separated specifically. A tree will try each feature once, and then select the one that can make the best feature of the classification as the parent node. The calculation of the best feature is based on the Gini coefficient, a splitting formula classic binary tree as below:

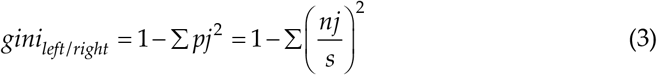

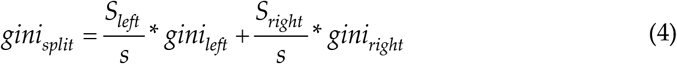

Where: Pj is the frequency column j appears in the total data set. Nj is the number of column j in the data set. S is the data numbers in the data set. The less Gini coefficient, the higher purity. In each split, the tree will choose the feature which the Gini is minimum as the split node.

A random forest is made up of 100 trees in the present research. A tree is made up of random number of samples and features. By voting result, it is decided by these 100 trees which category the data belongs to (the voting mechanism has one vote veto system, the minority subject to majority, and the weighted majority)

### 3.1.2. Density-Based Spatial Clustering of Applications with Noise (DBSCAN cluster)

DBSCAN is a density-based clustering algorithm. Such density clustering algorithms generally assume that categories can be determined by the closeness of the sample distribution. Samples of the same category are closely connected, which means there must be samples of the same category not far from any sample in the category. By classifying closely connected samples into one class, a clustering category is obtained. Classify all closely related samples into different categories, and the final results for all cluster categories are obtained. Two fundamental parameters of DBSCAN are EPS(ε) and Main points (MinPts). Eps is concept of the distance. For xj∈D, the ε-neighbor field contains the subsample set of the sample set D whose distance from xj is not greater than ε, Nε(xj)={xi ∈Distance(xi,xj)≤ε}, this sub The number of sample sets is recorded as |Nε(xj)|. For any sample xj∈D, if its ε-neighbor corresponding Nε(xj) contains at least MinPts samples, ie if |Nε(xj)|≥MinPts, then xj is the main points [1,8].

A benefit of utilizing DBSCAN is that the cluster is not based on a variety of distance metrics, but on density. Therefore, it can overcome the shortcomings of distance-based algorithms that can only find “circular-like” clusters. Meanwhile, the numbers of the cluster category cannot be set up in advance, which makes it much more objective than other clusters.

## 4. Result and Discussion

### 4.1. Current Land Type Classification and Soil Property

As is shown in the Table 2, 82 samples which comprise 4 current land types and are utilized to build the random forest classifier. After the establishment of the model, the gain of each feature is calculated and the other 18 samples will be classified by the model based on the gain of each feature sequentially.

**Table 2.**
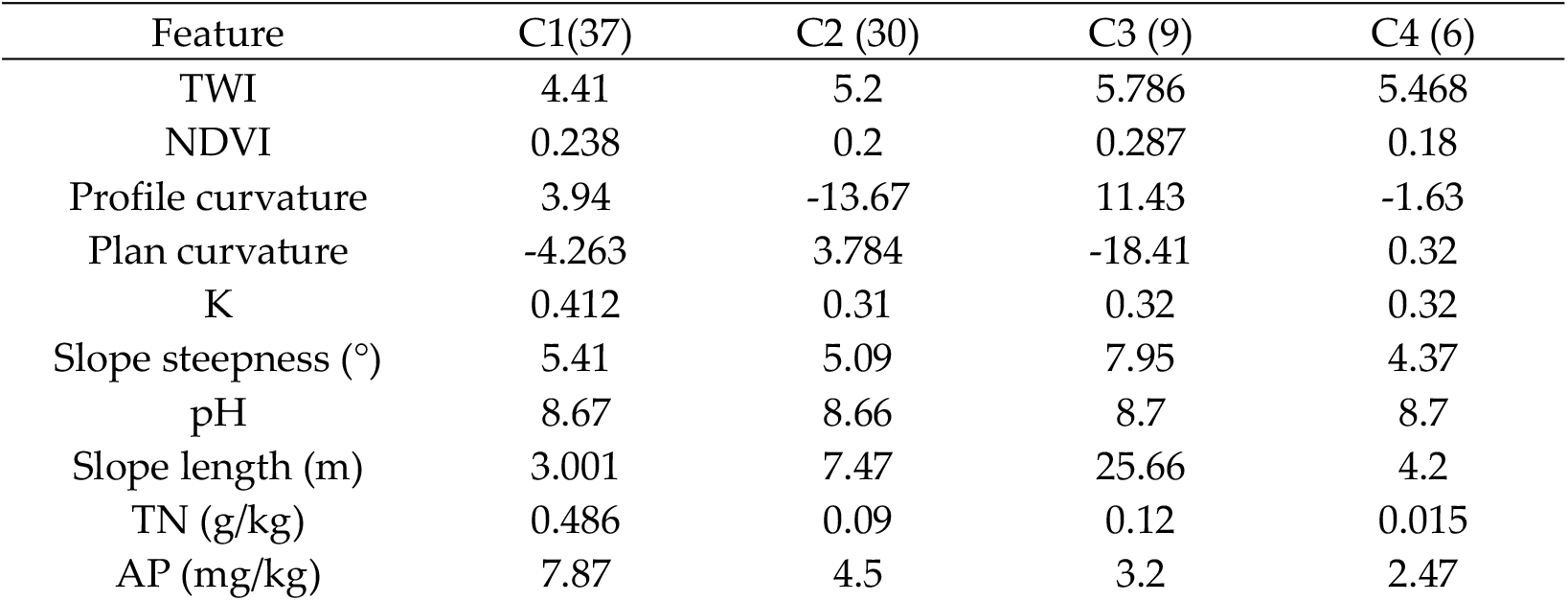
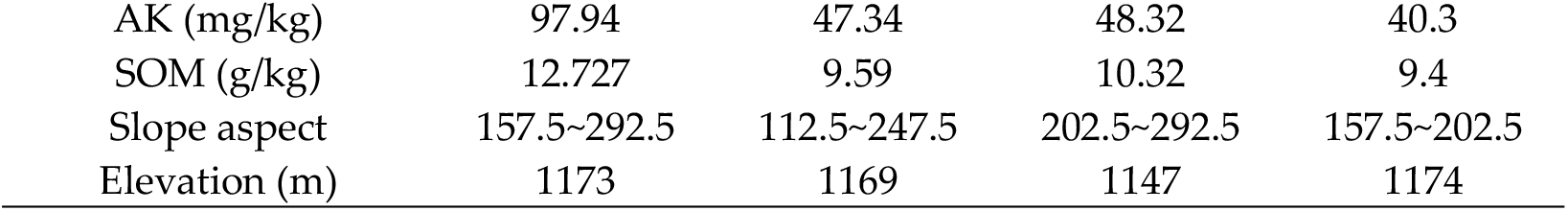
Mean value of environmental factors of the 100 samples.

The 82 samples comprise 32 cropland, 28 Shrub and grass land, 14 arbor land and 8 solar station. As is shown in the table, the soil nutrients comprise TN, AP, AK and SOM are most enriched in C1. The content of TN of C1 is 5 times more than C2 and 4 times more than C3. As is shown in the previous research, TN is the most significant nutrient affects the cropland [1]. Also, the K valve of C1 column is maximum and over 0.4, which means the soil particle is relatively smaller and the soil texture of it is sandy loam. The TWI of C1 is minimum, in consideration of a positive valve of profile curvature and a negative valve of plan curvature, this is presumably induced by the slope length as the cropland is mostly at the top of the slope the TWI could decrease from the bottom to the top of a slope due to the accumulation of the run off. There are some similarities between C2 and C3 column. TN, AK and SOM in C3 is slightly higher than in C2. In terms of AP, C2 exceeds a little. Also, the K valve of these two columns is similar, demonstrating a same type of soil---loamy sand. The curvature indicates that C3 has been suffered a runoff accumulation and acceleration., which ensures the abundant moisture that arbor requires. While the curvature of C2 could decrease and disperse the run-off, such circumstance of moisture is suitable for the drought enduring shrub and grass.

By contrasting C1, C2 and C3, it can be inferred that the limiting factor of classification for C1 column is TN and K as the murphy and corn requires TN as the top priority soil nutrient and loamy soil texture. As for the C2 and C3 column, the major classify factor could be NDVI and curvature. The NDVI of C3 is 0.087% higher than C2 and the curvature of C3 accelerate the runoff well the C2 resists instead. In terms of C4, the construction land, proved to be the minimum content of soil nutrients.

### 4.2. Random Forest Classification of Post Mining Land

The 82 samples engaged in the construction of the classification model based on the Gini. In each tree, 14 environmental factors and 82 samples are randomly selected and constructed. One of the trees is as below, where 52 samples and 6 features are selected. The Gini is calculated in each split till it reaches to zero, and the split will terminate. The max depth of the tree was set as 4 in case the model is overfitting.

As is shown in the Figure 2, class 1=C1, class2=C2, class3=C3, class4=C4. The first parent note which distinguish C1 and C2 is TN. The left branch continues to classify the C1 and C2 with the criteria of AK while the right branch classifies C1 and C3 with the criteria of TWI. When the depth goes to 3, the left branch of true path classifies C2 and C3 with the criteria of plan curvature and the right branch classifies C1 and C4 with the criteria of K. The left branch of the false path terminates and the right branch classifies C2 and C3 with the criteria of slope length.

**Figure 2.**
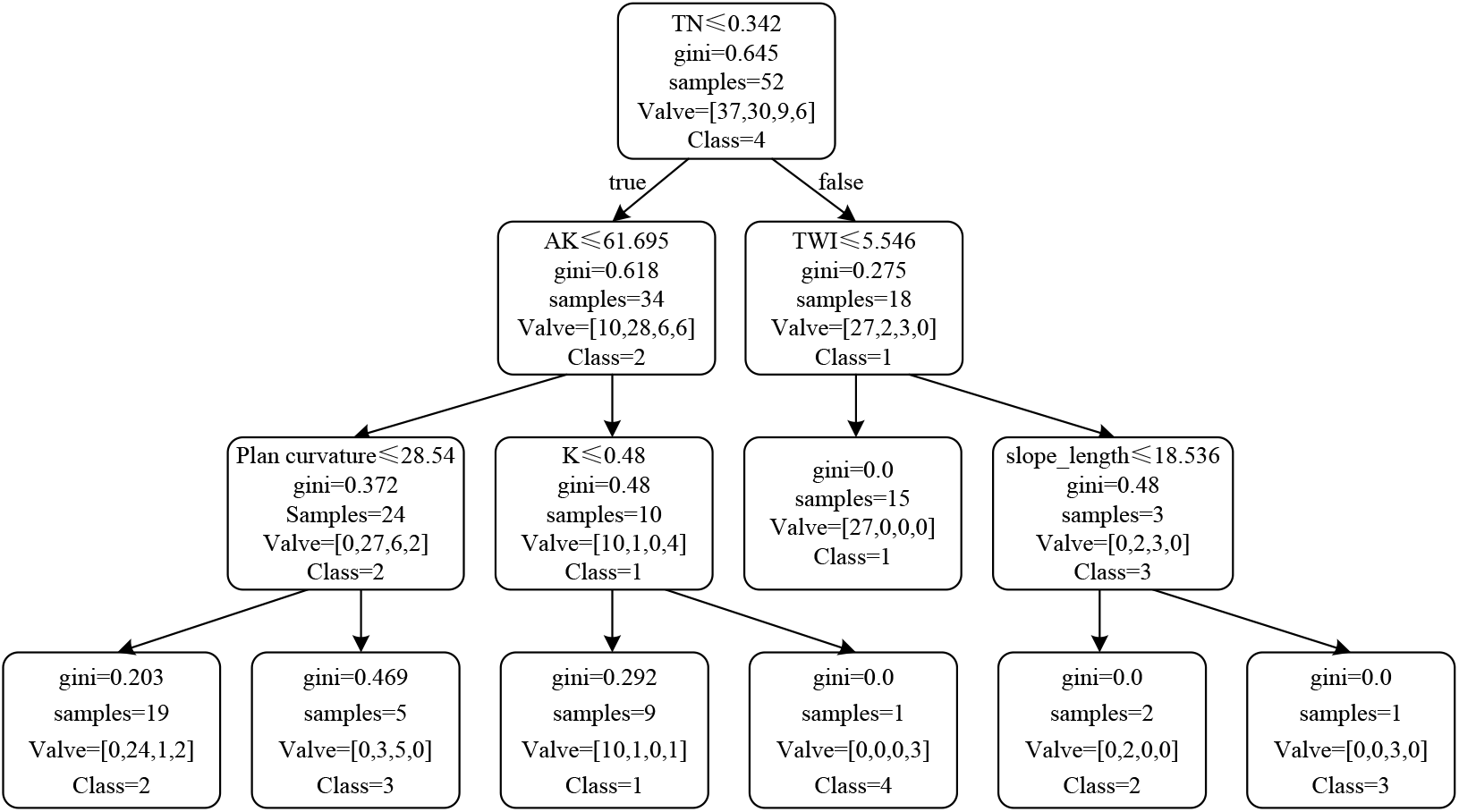
Classification tree build to predict the post-mining land use.

As the tree demonstrates, TN and AK are two major criteria distinguish C1 and C2, which indicates that theses two soil nutrients are with highly distinction in term of cropland and shrub land among the selected 6 features. While, the TWI is major criteria distinguish C1 and C3, indicating that a catchment area could conspicuously differentiate cropland and arbor land as the GINI decreased to zero. In the third depth, the plan curvature which distinguish the shrub land and arbor land is not efficient as the second depth, as in each branch the C2 and C3 consists in. The K valve is selected to distinguish shrub land and solar station, while the slope length is selected to distinguish shrub land and arbor land, which is generally accord with the previous data analysis. From the classifier tree, it can be inferred that C3 is literally distributed in the middle to the bottom of the slope and with high valve of TWI. C2 is basically distributed in the upper and middle of the slope.

### 4.3. Classification for the Test and Train Index

18 samples are utilized to examine the precision of the classification model as validation data-set. As is shown in the Table 3 below, the accuracy of the prediction model is about 95%, which is perceived as an available model.

**Table 3.** Validation of the post-mining land use classification.

| Samples Number | Reality result | RF prediction |
| --- | --- | --- |
| 11 | C4 | C4 |
| 19 | C2 | C2 |
| 25 | C2 | C2 |
| 32 | C2 | C2 |
| 42 | C2 | C2 |
| 45 | C1 | C1 |
| 47 | C3 | C2 |
| 49 | C4 | C4 |
| 55 | C2 | C1 |
| 66 | C4 | C4 |
| 67 | C2 | C2 |
| 73 | C1 | C1 |
| 75 | C1 | C1 |
| 83 | C2 | C2 |
| 84 | C2 | C2 |
| 85 | C1 | C1 |
| 91 | C1 | C1 |
| 98 | C1 | C1 |

No.55 is the exclusive sample RF prediction is error, where the classifier mistakenly assigned C3 to C1. However, the soil property of No.55 is ambiguous for the classifier as below: TWI=5.711, NDVI=0.286, profile curvature=7.97, plan curvature=-11.8391, K=0.42, Slope steepness= 4.6, pH=8.6, slope length=18.45, height=1213, TN=0.02, AP=2.75, AK=42.895, SOM=12.91, slope aspect=296.551. This sample is normally perceived as the noise point as the soil nutrients are basically above the average except the content of TN. The K valve is literally attributed to the C1 while the NDVI attributed to C2. The Gini coefficient which a purity criterion selected is incapable to classify this sample.

The 45 samples are totally idle post mining land and were predicted based on the classifier. As is shown in the Table 4, 15 samples were classified into C1, 23 samples were classified into C2, 5 samples were classified into C3, 2 samples were classified into C4. The valve of K and content of soil nutrients of the samples classified to be C1 column(C1-C) is maximum. Simultaneously, the slope steepness(°) is below 5° and is perceived as gentle surface, which is literally appropriate for reclaiming to cropland. C2-C, a relatively high content of soil available nutrients and the minimum valve of K, is suitable for the shrub. Considering the low valve of NDVI, the samples to be reclaimed to shrub and grass land could contribute to the ecological restoration by decreasing the soil erosion at the upper slope. Five samples are classified to the C3-C and the slope length of which is maximum while the content of TN and SOM is relatively high. which is accord with the circumstance that the site afforestation requires, the sunny slope could also ensure the sunshine and temperature condition for arbor. The two samples which possess the minimum content of soil nutrients and slope steepness is classified to C4-C. A relatively flat and smooth surface and a sunny slope is suitable for constructing the solar station.

**Table 4.** Post-mining land use prediction and the responding environmental factors.

|  | C1-C(15) | C2-C(23) | C3-C(5) | C4-C(2) |
| --- | --- | --- | --- | --- |
| TWI | 4.2 | 5.1 | **5.93** | 5.4 |
| K | **0.41** | 0.18 | 0.24 | 0.34 |
| TN (g/kg) | **0.28** | 0.038 | 0.056 | 0.027 |
| AP (mg/kg) | **5.55** | 3.67 | 2.2 | 1.5 |
| AK (mg/kg) | **97.75** | 42.477 | 39.499 | 29.648 |
| SOM(g/kg) | **13.988** | 8.79 | 9.93 | 8.31 |
| Slope steepness (°) | 4.32 | 5.57 | 4.14 | **2.18** |
| Slope length (m) | 3.31 | 7.62 | **19.36** | 1.25 |
| NDVI | 0.18 | 0.13 | 0.18 | 0.11 |
| Profile curvature | 3.57 | **-5.05** | 0.2126 | 0.16 |
| Plane curvature | 2.006 | **6.2** | -0.012 | -0.01 |

By the application of the Random forest, 45 samples were classified as 15 C1, 23 C2, 5 C3 and 2 C4. The proportion of the classification does not match to the initial proportion (37 C1, 30 C2, 9 C3 and 6 C4), indicating that the original proportion is not the fundamental of the classification. Unlikely to the neural network and SVM, the Gini of the random forest increased the accuracy and objectivity.

### 4.4. DBSCAN Cluster Utilization to the C1 and C2

Among the 145 samples, 58 C1, 61 C2 is inclusive. A DBSCAN cluster is utilized to analyze the coupling between soil property and the terrain distribution so that the plants and crops to be reclaimed or the quality and suitability of the same cluster could be determined. For C1, the cluster index comprises TN, AP and SOM which indicates the fertility of the corn and murphy requires. When EPS is set as 3, C1 is separated into two clusters, Cluster 1 comprises 24 samples and the average content of TN, AP and SOM is 0.566g/kg, 11.93mg/kg and 19.975g/kg respectively, while Cluster 2 comprises 34 samples and the average content of TN, AP and SOM is 0.304g/kg, 3.12mg/kg and 8.36g/kg respectively.

As is shown in the Figure 3, x, y and z refers to the TN, AP and SOM. The green points (Cluster 1) encircle the orange points (Cluster 2), The result of the cluster could also be proved by the content of the AK in which the content of C1 is average 112mg/kg while C2 is 81mg/kg. It can be inferred that the Cluster 1 and Cluster 2 of C1 are two different scale of soil fertility. Generally, Cluster 1 is suitable for cultivating the corn and murphy or soybean while slightly lack of TN and urea could be applied under certain circumstances. In terms of Cluster 2, the soil nutrients are comparatively barren in which manure and aquasorb is required. Ammonium phosphate and farmyard manure could be utilized to ameliorate the soil fertility.

**Figure 3.**
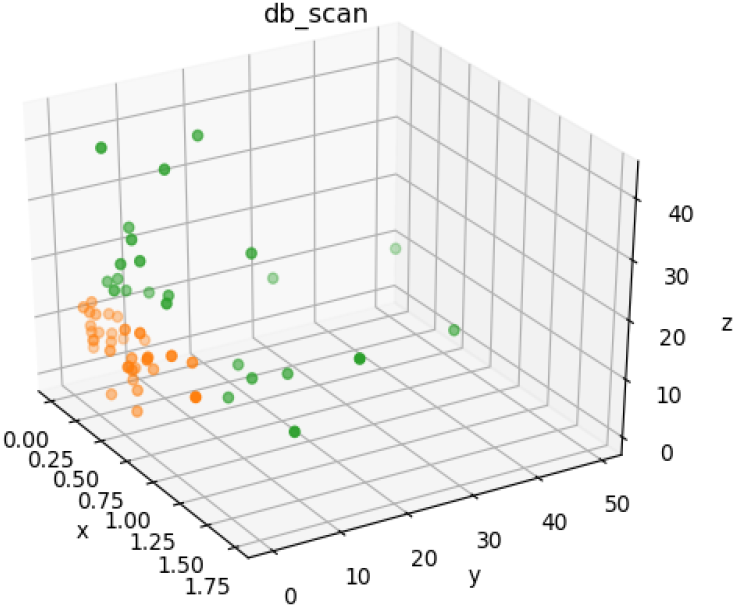
DBSCAN clustering based on SOM, TN and AP for C1.

As is shown in the Figure 4, the orange points (cluster 2) are basically distributed in the mined-out subsidence area and green points (cluster 1) are basically distributed in the non-collapsed area and valley, which evidently demonstrates that ground fracture and collapse pit has demolished the soil nutrients. In this situation, ultra-high-water material filling could be applied: Firstly, some coal gangue and loess with smaller particle size are used for deep filling. When the bottom crack is closed, the ultra-high-water material (bauxite and gypsum) is used for filling, of which 90% the content is water. After fully mixing the ground crack, the material quickly condenses within a few hours. At this time, the crack was filled with the soil nearby and the thickness was 50 cm. For Cluster 1, some corn production requires precision cultivation such as kx3564 could be planting. After filling the crack, the murphy and soybean could be planting in Cluster 2.

**Figure 4.**
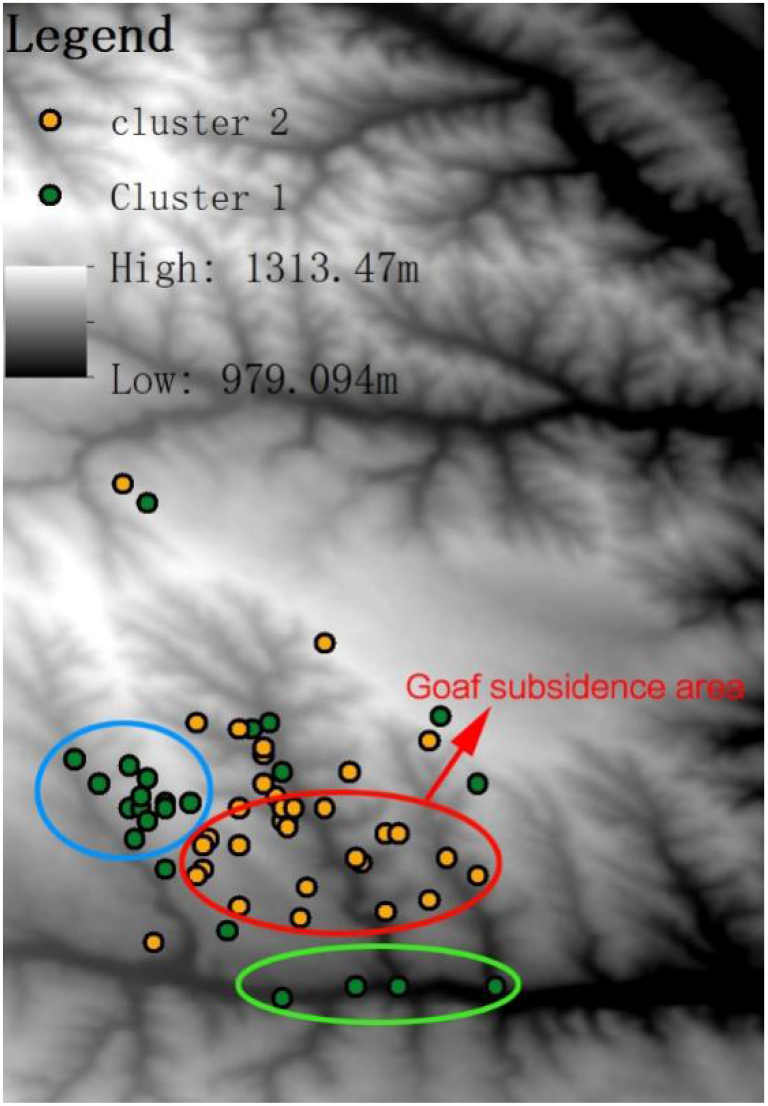
Spatial distribution of C1.

According to the previous research, AP and AK are the main nutrients C2 requires and slope aspect, slope steepness and K are three major factors limiting the site condition. As is analyzed, the slope steepness is generally below 10° and none of the slope is shady-slope or semi-shady slope. Therefore, the cluster index of C2 is selected as AP, AK and K. Data standardization is required as the unit of these three indexes are not uniform.

As is shown in the Figure 5, x, y and z refers to AP, AK and K. When EPS is set as 2, C2 is assembled by 3 parts. Cluster -1 comprises 24 samples and the average valve of AP, AK and K is 3.4mg/kg, 34.95mg/kg and 0.21. Cluster 0 comprise 22 samples and the average valve of AP, AK and K is 3.09mg/kg, 46.07mg/kg and 0.45. The criteria of orange points (Cluster -1) and blue points (Cluster 0) is K. Cluster -2 comprises 15 samples and average valve of AP, AK and K is 6.6mg/kg, 58.53mg/kg and 0.37 respectively, in which the content of AP and AK is maximum and SOM is minimum.

**Figure 5.**
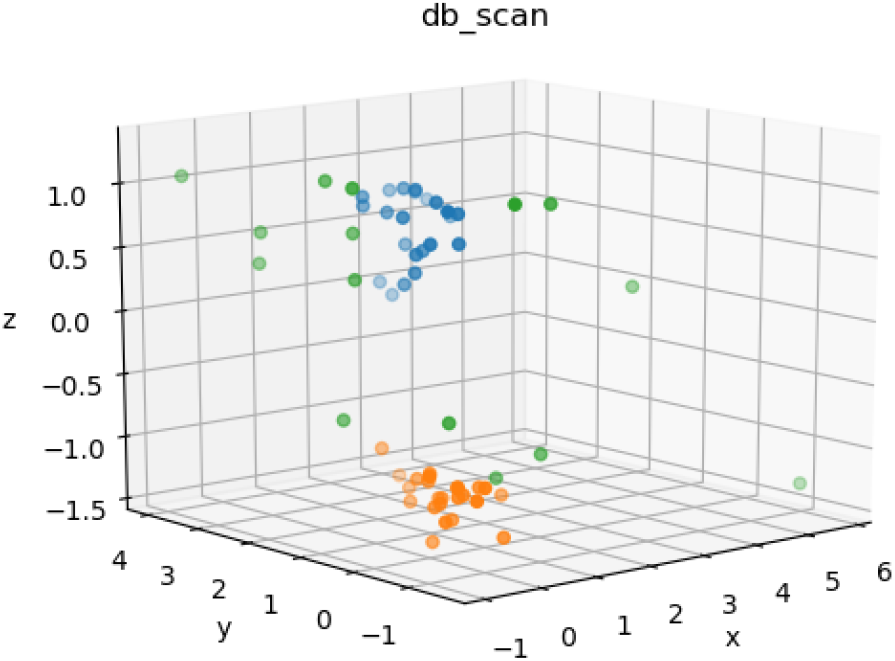
DBSCAN clustering based on AP and AK for C2.

As is shown in the Figure 6, Cluster −1 is basically distributed along the gully while Cluster 0 is distributed in the area with scarcely any gully. Such terrain distribution is presumably a reflection of the soil erosion consequence. In the gully area, large runoff induces the entrainment of small particles of the soil, and the large particles of soil left at the Cluster −1 result in a lower K value. Simultaneously, the runoff of Cluster 0 with barely gully is relatively small, so the small particles result in a higher K value. The AP values of Cluster −1 and 0 are similar. This is induced by the relatively low availability of phosphorus and both two Clusters are therefore less affected by water erosion. Based on the terrain distribution, it can be inferred that Cluster 0 is more suitable for planting grass plants, while Cluster 1 is more suitable for shrubs with higher plant height and bushes. grass requires small soil particles and soil aggregates with lower intensity of runoff. While shrubs such as caragana microphylla and Artemisia scoparia fit the larger soil particles.

**Figure 6.**
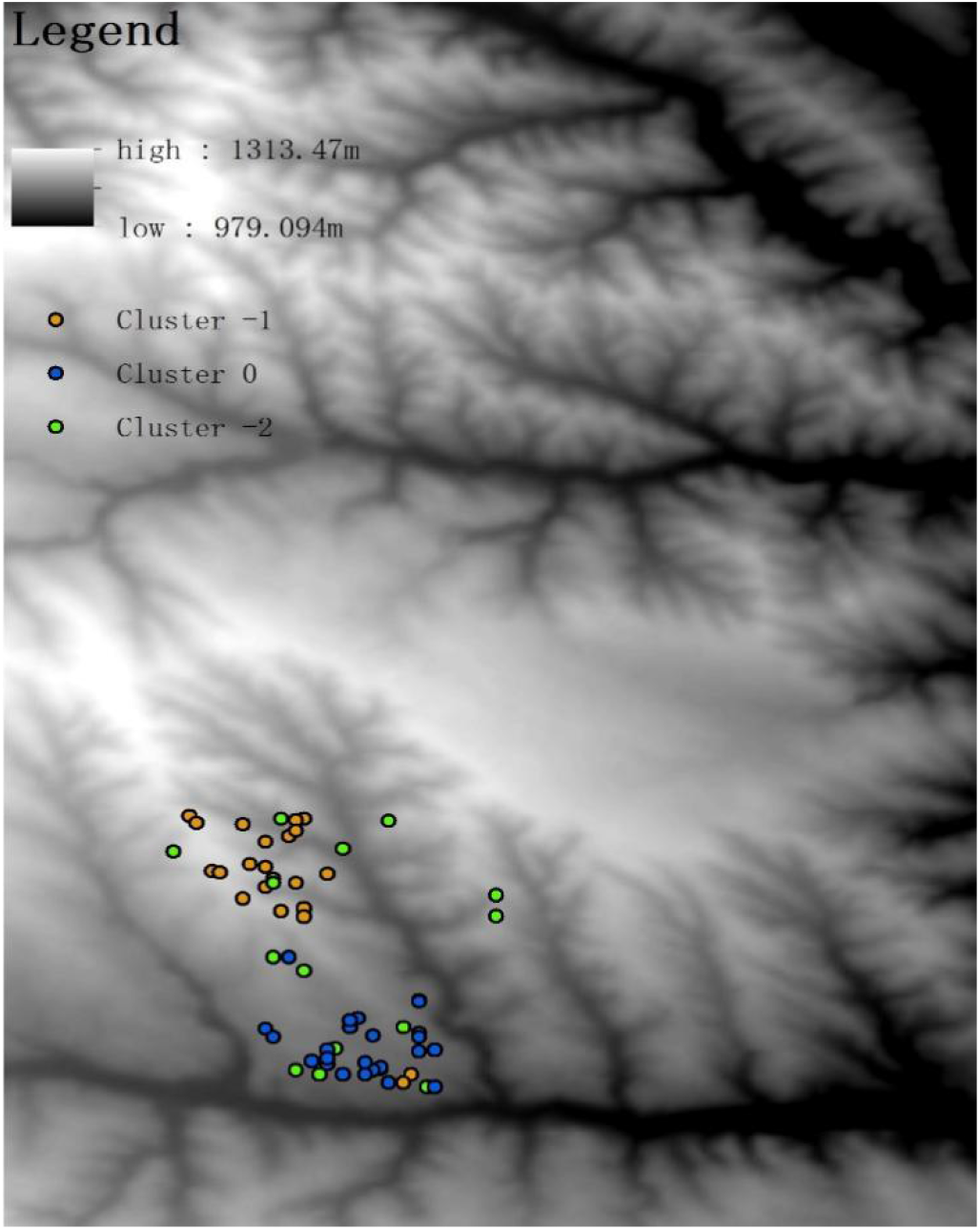
Spatial distribution of C2.

The distribution of Cluster 2 is generally scattered. The abundant soil available nutrients are ideal for leguminous plants. As is demonstrated in the previous research, the leguminous such as alfalfa and sweet clover requires abundant AP and AK to germinate. With its grow, it can gradually increase the content of SOM and be utilized into another shrub or arbor land.

## 5. Conclusion

In this study, 145 soil samples contains 45 to be reclaimed and 100 existing classification land use types were collected in order to build the post-mining land use classification model. Among the 100 samples, 82 samples were utilized to build the random forest classifier. The content of TN of C1 is 5 times more than C2 and 4 times more than C3. Also, the K valve of C1 column is maximum and over 0.4, which means the soil particle is relatively smaller and the soil texture of it is sandy loam. The TWI of C1 is minimum, in consideration of a positive valve of profile curvature and a negative valve of plan curvature, this is presumably induced by the slope length as the cropland is mostly at the top of the slope the TWI could decrease from the bottom to the top of a slope due to the accumulation of the run off. The accuracy of the prediction model is about 95%, which is perceived as an available model. In terms of the 45 to be reclaimed samples, 15 samples were classified into C1, 23 samples were classified into C2, 5 samples were classified into C3, 2 samples were classified into C4. The valve of K and content of soil nutrients of the samples classified to be C1 column(C1-C) is maximum. Simultaneously, the slope steepness(°) is below 5° and is perceived as gentle surface, which is literally appropriate for reclaiming to cropland. Total of 58 samples of C1 were clustered based on SOM, TN and AP. Cluster 1 comprises 24 samples and the average content of TN, AP and SOM is 0.566g/kg, 11.93mg/kg and 19.975g/kg respectively, while Cluster 2 comprises 34 samples and the average content of TN, AP and SOM is 0.304g/kg, 3.12mg/kg and 8.36g/kg respectively. Cluster 1 is suitable for cultivating the corn and murphy or soybean while slightly lack of TN and urea could be applied under certain circumstances. In terms of Cluster 2, the soil nutrients are comparatively barren in which manure and aquasorb is required. Ammonium phosphate and farmyard manure could be utilized to ameliorate the soil fertility.

## Author Contributions

Conceptualization, Qiang Bo and Pinhan Lv.; methodology, Zechuan Li.; software, Qian Wang.; validation,Ziguan Wang.; writing—original draft preparation, Qiang Bo.; writing—review and editing, Zi guan wang.

## Acknowledgments

The work represents a contribution to the foundation of Beijing brilliant youth experts. We thank Qingyu Xu, Yunfei Bai for their kindly help.

## Disclaimer/Publisher’s Note

The statements, opinions and data contained in all publications are solely those of the individual author(s) and contributor(s) and not of MDPI and/or the editor(s). MDPI and/or the editor(s) disclaim responsibility for any injury to people or property resulting from any ideas, methods, instructions or products referred to in the content.

